# The PedS2/PedR2 two-component system is crucial for the rare earth element switch in *Pseudomonas putida* KT2440

**DOI:** 10.1101/364661

**Authors:** Matthias Wehrmann, Charlotte Berthelot, Patrick Billard, Janosch Klebensberger

**Affiliations:** University of Stuttgart, Institute of Technical Biochemistry, Stuttgart, Germany; Université de Lorraine, LIEC UMR7360, Faculté des Sciences et Technologies, Vandoeuvre-lès-Nancy, France; CNRS, LIEC UMR7360, Faculté des Sciences et Technologies, Vandoeuvre-lès-Nancy, France

**Keywords:** Lanthanides, rare earth element switch, signal transduction, *Pseudomonas putida*, PQQ, two-component regulatory system, PedS2, PedR2, histidine kinase, LuxR-type regulator, dehydrogenases, periplasm

## Abstract

In *Pseudomonas putida* KT2440, two pyrroloquinoline quinone-dependent ethanol dehydrogenases (PQQ-EDHs) are responsible for the periplasmic oxidation of a broad variety of volatile organic compounds (VOCs). Depending on the availability of rare earth elements (REEs) of the lanthanide series (Ln^3+^), we have recently described that the transcription of the genes encoding the Ca^2+^-utilizing enzyme PedE and the Ln^3+^-utilizing enzyme PedH are inversely regulated. With adaptive evolution experiments, site-specific mutations, transcriptional reporter fusions, and complementation approaches, we herein demonstrate that the PedS2/PedR2 (PP_2671/PP_2672) two-component system (TCS) plays a central role in the observed REE-mediated switch of PQQ-EDHs in *P. putida*. We provide evidence that in the absence of lanthanum (La^3+^), the sensor histidine kinase PedS2 phosphorylates its cognate LuxR-type response regulator PedR2, which in turn not only activates *pedE* gene transcription but is also involved in repression of *pedH*. Our data further suggests that the presence of La^3+^ lowers kinase activity of PedS2, either by the direct binding of the metal ions to the periplasmic region of PedS2 or by an uncharacterized indirect interaction, leading to reduced levels of phosphorylated PedR2. Consequently, the fading *pedE* expression and concomitant alleviation of *pedH* repression causes – in conjunction with the transcriptional activation of the *pedH* gene by a yet unknown regulatory module – the Ln^3+^-dependent transition from PedE to PedH catalysed oxidation of alcoholic VOCs.

**IMPORTANCE:** The function of lanthanides for methano- and methylotrophic bacteria is gaining increasing attention, while knowledge about the role of rare earth elements (REEs) in non-methylotrophic bacteria is still limited. The present study investigates the recently described differential expression of the two PQQ-EDHs of *P. putida* in response to lanthanides. We demonstrate that a specific TCS is crucial for their inverse regulation and provide evidence for a dual regulatory function of the LuxR-type response regulator involved. Thus, our study represents the first detailed characterization of the molecular mechanism underlying the REE switch of PQQ-EDHs in a non-methylotrophic bacterium and stimulates subsequent investigations for the identification of additional genes or phenotypic traits that might be co-regulated during REE-dependent niche adaptation.

## INTRODUCTION

In its natural soil habitat, *Pseudomonas putida* KT2440 is exposed to a broad range of potential carbon and energy sources (Regenhardt et al., 2002; Wackett, 2003; Wu et al., 2011), including plant-, fungi-, or bacterially derived volatile organic compounds (VOCs) with alcohols or aldehydes as functional groups (Insam and Seewald, 2010; Penuelas et al., 2014; van Dam et al., 2016). For efficient capture and metabolism of such VOCs, *P. putida* makes use of two PQQ-dependent ethanol dehydrogenases (PQQ-EDHs) – namely PedE and PedH – to carry out the initial oxidation steps in the periplasm of the cell (Mückschel et al., 2012; Wehrmann et al., 2017). In a recent study, we found that these two type I quinoproteins (Anthony, 2001; Toyama et al., 2004) exhibit a similar substrate scope, but require different metal cofactors for their catalytic activity (Wehrmann et al., 2017). In contrast to PedE, which uses Ca^2+^ ions, PedH was characterized as a rare earth element (REE)-dependent enzyme that relies on the presence of lanthanide (Ln^3+^) ions for catalytic activity. Notably, due to their low solubility in most natural environments, REEs have long been considered to have no biological function (Sarkar, 2002). However, the discovery of the wide-spread occurrence of the REE-dependent XoxF-type of PQQ-dependent methanol dehydrogenases (PQQ-MDHs), together with the more recent description of Ln^3+^-dependent ethanol dehydrogenases, has highlighted the important role of REEs for many bacterial species in various environmental compartments (Fitriyanto *et al*., 2011a; Fushimi, et al., 2011; Pol et al., 2014; Taubert et al., 2015; Chu and Lidstrom, 2016; Good et al., 2016; Vekeman et al., 2016; Shiller et al., 2017; Wehrmann et al., 2017; Lv et al., 2018; Masuda et al., 2018). While in the absence of Ln^3+^ the oxidation of methanol in methylotrophs is driven by Ca^2+^-dependent PQQ-MDHs (MxaF-type), the presence of low amounts of REE ions is usually sufficient to trigger a transcriptional switch to the XoxF-type of PQQ-MDHs. This inverse regulation, called the REE switch, has been described for many methano- and/or methylotrophic organisms (Farhan Ul Haque et al., 2015; Chu and Lidstrom, 2016; Vu et al., 2016; Gu and Semrau, 2017; Masuda et al., 2018; Zheng et al., 2018). From the growing number of studies, it has become apparent that the molecular mechanism underlying this switch for PQQ-MDHs is complex and can substantially differ among species. For example, the inverse regulation in the non-methanotrophic methylotroph *Methylobacterium extorquens* strain AM1 is controlled by two different two-component systems (TCS) (MxcQE and MxbDM) and the orphan response regulator MxaB (Xu et al., 1995; Springer et al., 1997, 1998). In this organism it has been found that the transcriptional activation of both enzymes, the Ca^2+^-dependent MxaF and the two Ln^3+^-dependent XoxF1 and XoxF2, is entirely lost in a Δ*xoxF1*Δ*xoxF2* double mutant (Skovran et al., 2011). As a consequence, a complex and consecutive regulation was postulated, in which the different binding affinities of the *apo* (no Ln^3+^ bound to the enzyme) and *holo* (Ln^3+^ bound to the enzyme) form of the XoxF p roteins to the periplasmic domain of the sensor histidine kinase MxcQ is essential to regulate the switch (Vu et al., 2016).

The type I methanotroph *Methylomicrobium buryatense* strain 5GB1C is lacking homologs of the aforementioned TCS systems MxcQE and MxbDM (Chu and Lidstrom, 2016). In this organism, the REE switch is regulated predominantly by the sensor histidine kinase MxaY (Chu et al., 2016). Chu and coworkers found that the activation of MxaY in the absence of lanthanum activates the transcription of the Ca^2+^-dependent enzyme MxaF by a so-far unknown response regulator. In addition, the deletion of MxaY was found to almost entirely eliminate the Ln^3+^-mediated transcriptional activation of the Ln^3+^-dependent enzyme XoxF in a partially MxaB-dependent manner that also results in a severe growth defect, both in the absence and presence of Ln^3+^. As an additional layer of complexity, recent studies found that the presence of other metal ions such as copper, which is needed as a cofactor for the membrane-bound or particulate methane monooxygenase (pMMO) in methanotrophs, can significantly impact the REE-mediated switch (Farhan Ul Haque et al., 2015; Gu and Semrau, 2017; Semrau et al., 2018).

In contrast to the increasing knowledge about the regulation of PQQ-MDHs in methylotrophs, the molecular basis underlying such a regulatory switch for PQQ-EDHs in non-methylotrophic organisms is not established. In the present study, we identify the TCS encoded by PP_2671/PP_2672 (hereinafter referred to as PedS2/PedR2 according to the genetic nomenclature from Arias et al., 2008), consisting of the sensor histidine kinase PedS2 and its cognate LuxR-type transcriptional response regulator PedR2, as an essential regulatory module for the REE-mediated switch of PQQ-EDHs in *P. putida* KT2440. We provide evidence that the activity of PedS2 in the absence of lanthanides leads to phosphorylation of PedR2 (PedR2^P^), which serves a dual regulatory function. On the one hand, PedR2^P^ acts as a strong transcriptional activator of the *pedE* gene, which is essential to allow growth with 2-phenylethanol in the absence of Ln^3+^. At the same time, PedR2^P^ also functions as a repressor of *pedH*. From our data, we conclude that the presence of Ln^3+^ ions triggers a reduction in PedS2 activity, either by a direct binding of the metal to the periplasmic region of PedS2 or by an uncharacterized indirect interaction. This reduction of PedS2 activity, together with a proposed phosphatase activity of PedS2 in its inactivated state, causes the accumulation of non-phosphorylated PedR2, which over-time results in the loss of the regulatory activity of the protein, and facilitates – in concert with the positive feedback function of PedH in the presence of Ln^3+^ ions – the switch between PedE and PedH-dependent growth.

## RESULTS

### Identification of the sensor histidine kinase PedS2 as a lanthanide-responsive sensor

As a consequence of the inverse regulation of *pedE* and *pedH*, a *pedH* deletion strain does not grow within 48 h with 2-phenylethanol as sole carbon source in the presence of a critical concentration of La^3+^ in the culture medium (Wehrmann et al., 2017). To test whether strains can be evolved, which are capable of overcoming the repression of *pedE* in the presence of Ln^3+^ ions, an adaptive evolution experiment was performed (see **Figure 1** for a general scheme).

**Figure 1:**
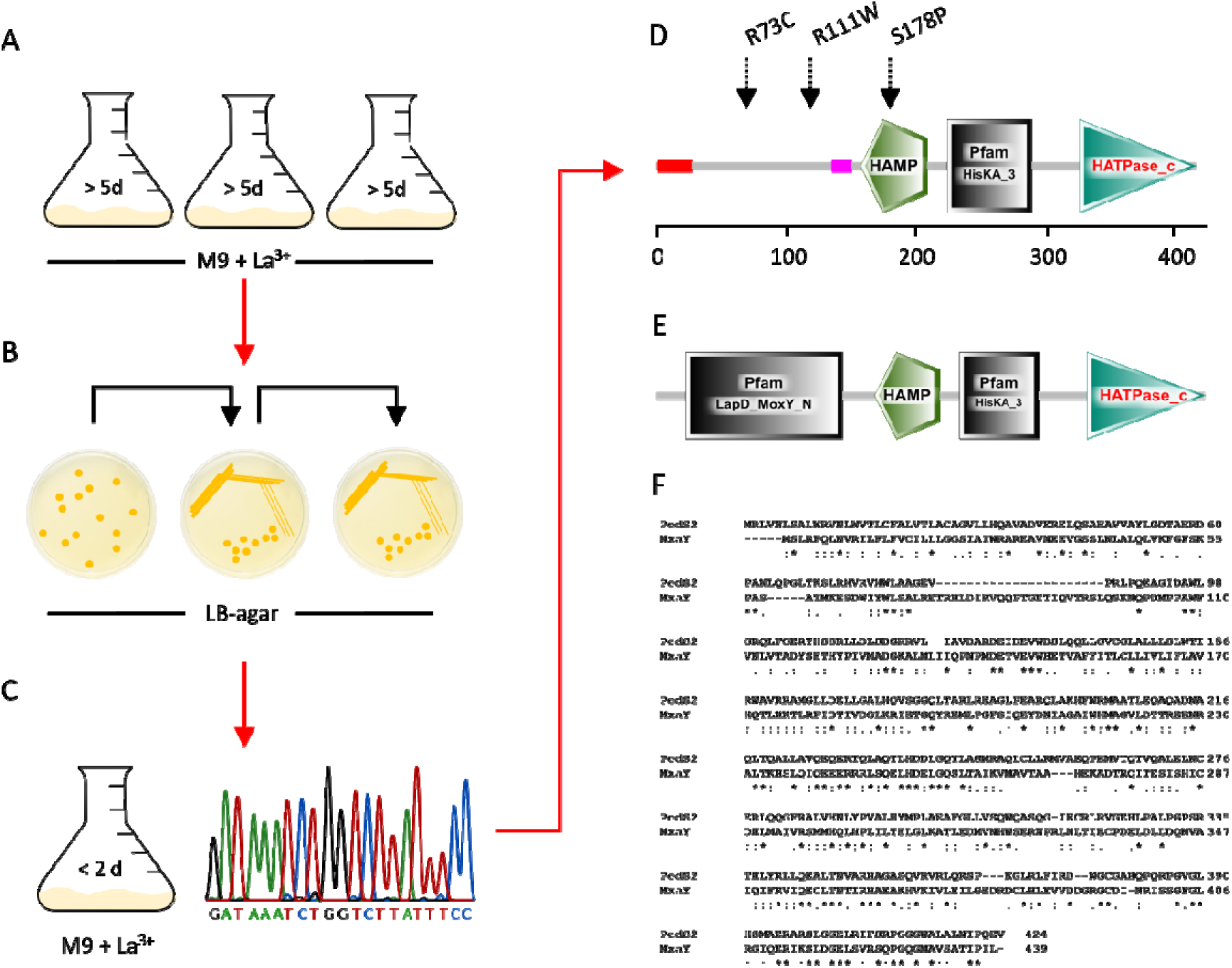
Scheme of selection (**A**), clonal isolation (**B**), characterization and single nucleotide polymorphism (SNP) identification (**C-D**) in the two-component sensor histidine kinase PedS2 of the spontaneous mutants ∆*pedH^S^*^1^ (R73C), ∆*pedH^S^*^2^ (R111W), ∆*pedH^S^*^3^ (S178P). A) Cells of strain ∆*pedH* were incubated in M9 medium supplemented with 5 mM 2-phenylethanol and 10 µM LaCl_3_ in plastic Erlenmeyer flasks (n = 3) at 30°C with shaking at 180 rpm. **B**) After growth was observed (> 5 days), dilutions from each culture were plated onto LB-agar plates and incubated at 30°C. Individual clones were further streaked on LB-agar twice prior to further characterization. C) Clones were characterized for their growth behavior in M9 medium with 5 mM 2-phenylethanol in the presence of 10 µM LaCl_3_. Subsequently, one clone from each culture exhibiting faster growth than the parental strain ∆*pedH* was used for PCR amplification of the *pedS2* gene and multi sequence alignment analysis with the native sequence of the gene from the *Pseudomonas* Genome Database (Winsor et al., 2016). **D-E**) Visualization of domain composition of PedS2 of *P. putida* (**D**) and MxaY of *M. buryatense* 5GB1C (**E**) using the prediction from the Simple Modular Architecture Research Tool (Letunic and Bork, 2018). **F**) Amino acid sequence alignment of the PedS2 and MxaY proteins generated with Clustal Omega (Sievers et al., 2014).

When Δ*pedH* cultures were incubated with 2-phenylethanol in the presence of 10 µM La^3+^ ions for longer than 5 d, growth was observed, indicating the occurrence of adapted strains (data not shown). When independent clones were isolated from such cultures and passaged several times on LB agar medium, their growth phenotype with 2-phenylethanol was still much faster (< 2 d) than that observed for their Δ*pedH* parental strain and very similar to the growth phenotype of the wild-type strain KT2440. Similar spontaneous suppressor mutants have been described recently in methylotrophic organisms such as *M. extorquens* strain AM1, *M. buryatense strain* 5GB1C, and *M. aquaticum* strain 22A during growth with methanol (Chu and Lidstrom, 2016; Chu et al., 2016; Masuda et al., 2018). In *M. buryatense*, whole genome sequencing revealed that the suppressor mutant strain was characterized by a mutation in the membrane bound two-component sensor histidine kinase MxaY (Chu et al., 2016). In *P. putida* KT2440, the gene PP_2671 (hereinafter referred to as *pedS2*), located in close genomic proximity to *pedE* (PP_2674), encodes a membrane bound histidine kinase sharing 25 % amino acid sequence identity with MxaY (**Figure 1D-F**). To test the hypothesis that mutations in *pedS2* are responsible for the emergence of suppressor phenotypes in strain ∆*pedH* during growth in the presence of La^3+^, the gene was sequenced in three individually isolated mutants (∆*pedH^S^*^1-3^) (**Figure 1A-D**). This analysis revealed that all strains contained a mutation in the *pedS2* sequence leading to a non-synonymous substitution. These mutations were either located within a predicted periplasmic domain of unknown function (R73C [∆*pedH^S^*^1^] and R111W [∆*pedH^S^*^2^]) or within the HAMP domain (S178P [∆*pedH^S^*^3^]), which is responsible for signal transduction from the periplasm into the cytoplasm of the cell (Stock et al., 2000; Parkinson, 2010). To verify that the identified mutations in *pedS2* are the primary cause of the suppressor phenotype, the S178P mutation from ∆*pedH^S^*^3^ was introduced into the genetic background of strain ∆*pedH*, resulting in strain ∆*pedH_*PedS^S178P^.

In subsequent growth experiments with 2-phenylethanol in the absence of La^3+^, strains ∆*pedH* and ∆*pedH_*PedS2^S178P^ showed a similar growth behavior with a lag phase of < 32 h (**Figure 2A1; Figure 2A2**). In the presence of 10 µM La^3+^, however, ∆*pedH* showed no growth within 72 h, whereas ∆*pedH_*PedS2^S178P^ reached its maximum optical densities again after about 32 h of incubation, verifying that the observed mutation in the histidine kinase *pedS2* was sufficient to cause the suppressor phenotype. We speculated that the LuxR-type response regulator PP_2672 (hereinafter referred to as *pedR2*), which is located adjacent to *pedS2* within the genome of *P. putida* KT2440, represents the target of PedS2 activity. This assumption is based on the fact that PedR2 represents a homologue (65% amino acid sequence identity) of EraR (ExaE; PA1980), which forms a TCS with the cytosolic histidine kinase EraS (ExaD; PA1979) that activates expression of the *pedE* homologue *exaA* in *Pseudomonas aeruginosa* (Schobert and Görisch, 2001; Mern et al., 2010). To test this hypothesis, *pedS2* as well as its potential cognate response regulator encoding gene *pedR2*, were deleted in a ∆*pedH* background. In addition, strains suitable for probing promoter activity of *pedE* in strains ∆*pedH*, ∆*pedH_*PedS2^S178P^, ∆*pedH*∆*pedS2* and ∆*pedH*∆*pedR2* were constructed and subsequently analyzed during growth with 2-phenylethanol in the presence and absence of La^3+^ (**Figure 2B**).

**Figure 2:**
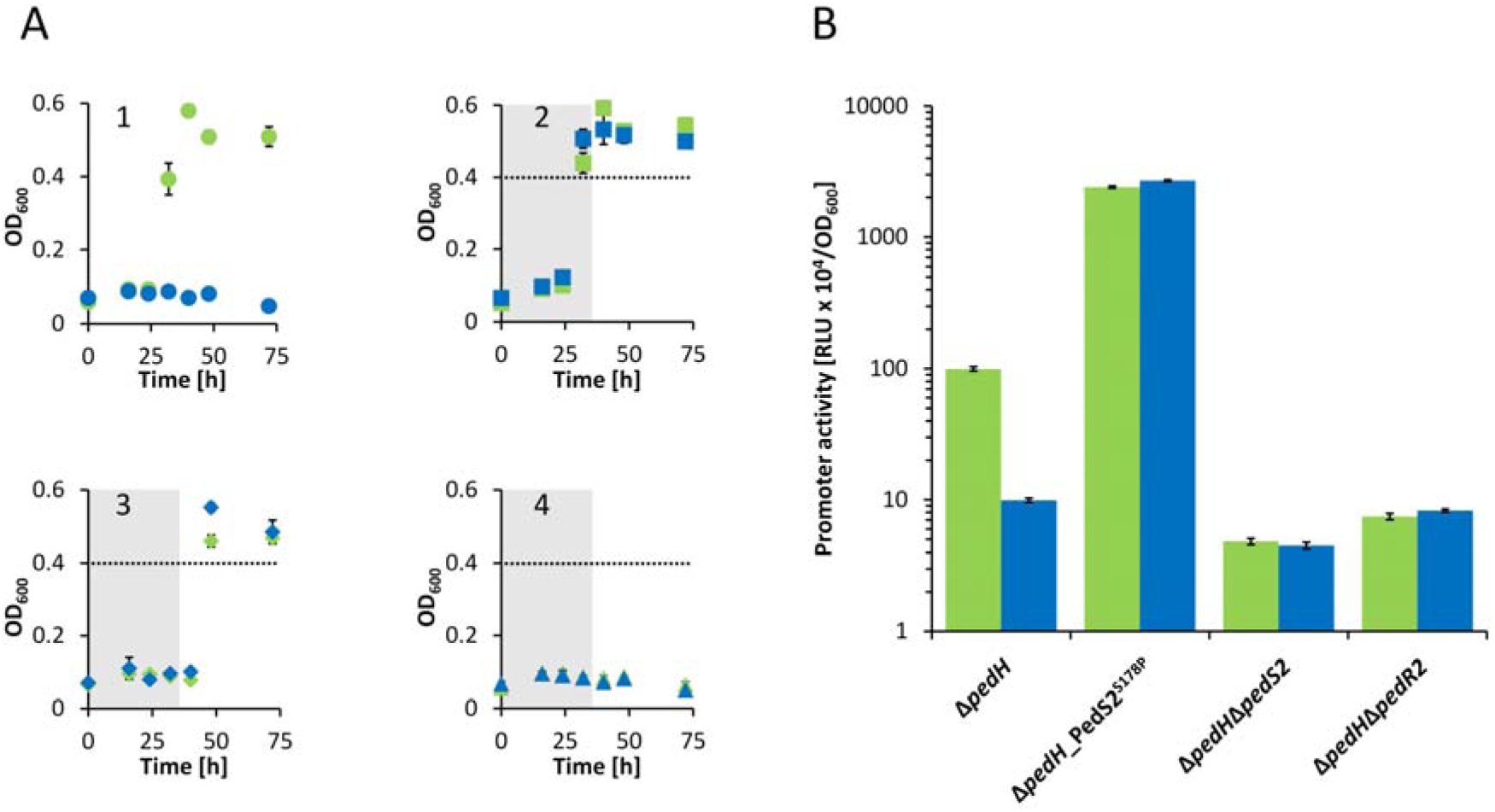
(**A**) Growth of strains ∆pedH (circles; 1), ∆*pedH*_*pedS2^S178P^* (squares; 2), ∆*pedH*∆*pedS2* (diamonds; 3), and ∆*pedH*∆*pedR2* (triangles; 4) at 30°C and 350 rpm shaking with M9 medium in 96-well plates supplemented with 5 mM 2-phenylethanol in absence (green symbols) or presence (blue symbols) of 10 µM La^3+^. The grey area visualizes the time point by which the parental strain ∆*pedH* (circles) reached an OD_600_ > 0.4 (dotted line) (**B**) Activities of the *pedE* promoter in strains ∆*pedH*, ∆*pedH*_PedS2^S178P^, ∆*pedH*∆*pedS2*, and ∆*pedH*∆*pedR2* in the absence (green bars) or presence (blue bars) of 1 µM La^3+^ measured in M9 medium supplemented with 1 mM 2-phenylethanol. Promoter activities are presented in relative light units (RLU x 10^4^) normalized to OD_600_. All data represent the mean of biological triplicates and error bars correspond to the respective standard deviations.

### The TCS PedS2/PedR2 regulates pedE transcription in response to lanthanide availability

In accordance with the observed growth patterns, *pedE* promoter activities in strain ∆*pedH_*PedS2^S178P^ were almost identical in both the absence and the presence of La^3+^ (ratio no La^3+^ to 1 µM La^3+^: 0.89 ± 0.02) but increased more than 24-fold (24 ± 1 and 27 ± 1-fold, respectively) compared to the *pedE* promoter activities determined for cells of strain ∆*pedH* in the absence of La^3+^ (**Figure 2B**). Strain ∆*pedH*∆*pedS2* also showed a La^3+^-independent growth phenotype similar to strain ∆*pedH_*PedS2^S178P^, however with a clear delay (< 48 h vs. < 32 h) (**Figure 2A3**). Promoter activities for *pedE* were almost identical in this strain in the absence and presence of La^3+^ (ratio no La^3+^ to 1 µM La^3+^ : 1.07 ± 0.09) (**Figure 2B**).

In contrast, incubations of 72 h (**Figure 2A4**) or even prolonged incubations for more than 7 d (data not shown) did not result in detectable growth of strain ∆*pedH*∆*pedR2* with 2-phenylethanol, both in the presence and absence of La^3+^. Correspondingly*, pedE* promoter activities in this strain were low compared to those observed for cells of strain ∆*pedH* in the presence of La^3+^ but in a similar range as those observed for ∆*pedH*∆*pedS2* in the presence and absence of La^3+^. These data demonstrate that the PedS2/PedR2 system is the predominant element in the La^3+^ -dependent regulation of *pedE* and that PedR2 is essential for PedE-dependent growth. However, as PedE-dependent growth can still be observed in the absence of PedS2 after prolonged incubations and in a PedR2-dependent manner (**Figure 2A3; Figure 2A4**), we assume that at least one additional lanthanide-independent kinase must be able to phosphorylate PedR2, leading to transcriptional activation of *pedE* and functional production of the calcium-dependent enzyme under these conditions.

### The TCS PedS2/PedR2 regulates the partial repression of pedH in the absence of lanthanides

Based on the critical role of PedS2/PedR2 in the regulation of *pedE* and the fact that LuxR-type regulators have been demonstrated to be capable of acting as both transcriptional activators and repressors (Waters and Bassler, 2006; Van Kessel et al., 2013), we speculated that this TCS could also be involved in the regulation of *pedH*. To test this hypothesis, strains ∆*pedE*_PedS2^S178P^, ∆*pedE*∆*pedS2*, and ∆*pedE*∆*pedR2* were generated and a transcriptional reporter suitable for probing *pedH* promoter activities was integrated into the genome of each of these strains.

Experiments with 2-phenylethanol revealed that strains ∆*pedE*,∆*pedE*∆*pedS2*, and ∆*pedE*∆*pedR2* showed La^3+^-dependent growth after a lag-phase < 24 h and reached the maximum OD_600_ after ≤ 32 h (**Figure 3A1**; **Figure 3A3**; **Figure 3A4**). In contrast, strain ∆*pedE*_PedS2^S178P^ exhibited an extended lag-phase (> 24 h) and consistently reached the maximal OD_600_ only after prolonged incubations (≥ 32 h; **Figure 3A2**). In accordance with these growth results, *pedH* promoter activities of strains ∆*pedE*, ∆*pedE*∆*pedS2*, and ∆*pedE*∆*pedR2* were in a similar range, whereas *pedH* promoter activities in strain ∆*pedE*_PedS2^S178P^ were 45 ± 2-fold and 30 ± 2-fold lower than strain ∆*pedE* in absence or presence of La^3+^, respectively (**Figure 3B**). Assuming that the S178P mutation in PedS2 results in a sensor kinase activity that mimics that of the wild-type protein in the absence of lanthanides, it is very likely that PedS2 is responsible for the repression of *pedH* under the La^3+^-free conditions leading to the observed delay in growth. To find out whether this regulatory effect on *pedH* proceeds *via* the response regulator PedR2, as it is the case for *pedE*, or is caused by an unknown additional target of PedS2, the triple mutant strain ∆*pedE*_PedS2^S178P^ ∆*pedR2* was generated and characterized for its growth phenotype (**Figure 4A**). In this experiment, ∆*pedE* and ∆*pedE*_PedS2^S178P^ both grew with 2-phenylethanol, however, with a clear difference in the corresponding lag-phases confirming the previous results from growth in 96-well plates (**Figure 3A**). In contrast, the additional deletion of the response regulator PedR2^S178P^ eliminated the growth defect caused by the PedS2 allele, leading to a growth behavior of ∆*pedE*_PedS2^S178P^ ∆*pedR2* (**Figure 4A**), which was indistinguishable from that of strain ∆*pedE*.

**Figure 3:**
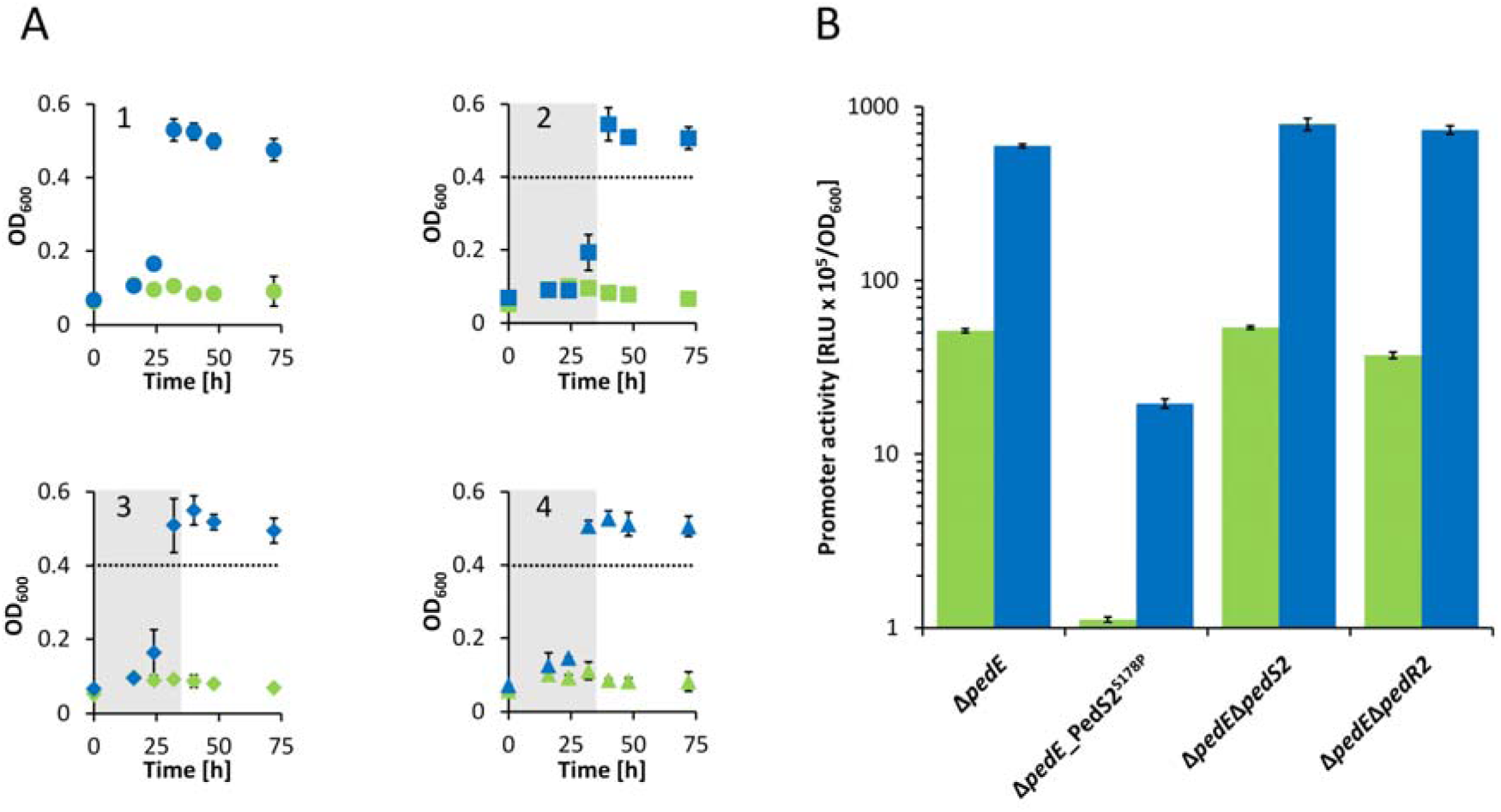
(**A**) Growth of strains Δ*pedE* (circles; 1), Δ*pedE*_PedS2^S178P^ (squares; 2), Δ*pedE*Δ*pedS2* (diamonds; 3), and Δ*pedE*Δ*pedR2* (triangles; 4) at 30°C and 350 rpm shaking with M9 medium in 96-well plates supplemented with 5 mM 2-phenylethanol in absence (green symbols) or presence (blue symbols) of 10 μM La^3+^424. The grey area visualizes the time point by which the parental strain Δ*pedE* (circles) reached their maximum OD_600_ (**B**) Activities of the *pedH* promoter in strains Δ*pedE*, Δ*pedE*_PedS2^S178P^, Δ*pedE*Δ*pedS2*, and Δ*pedE*Δ*pedR2* in the absence (green bars) or presence (blue bars) of 1 μM La^3+^ measured in M9 medium supplemented with 1 mM 2-phenylethanol. Promoter activities are presented in relative light u nits ( RLU x 10^5^) normalized to OD_600_. All data represent the mean of biological triplicates and error bars correspond to the respective standard deviations.

**Figure 4:**
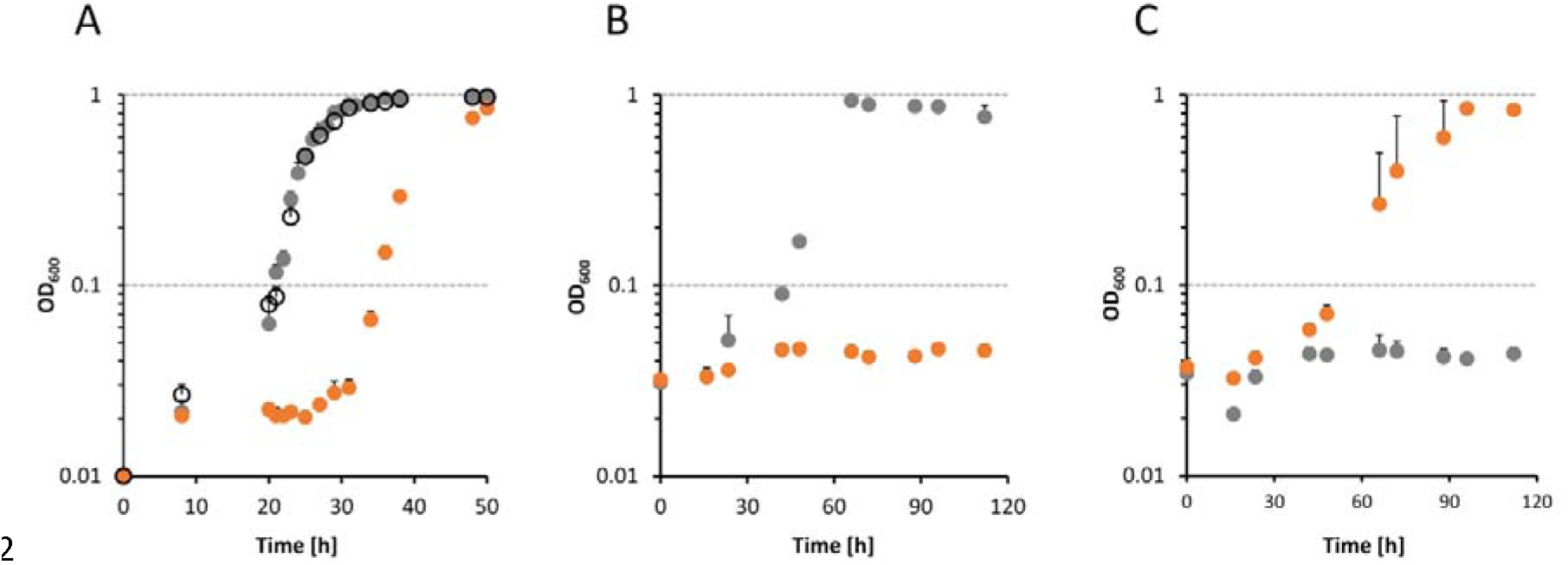
Growth of different *P. putida* strains at 30°C and 180 rpm shaking with M9 medium in polycarbonate Erlenmeyer flasks supplemented with 5 mM 2-phenylethanol and 10 μM La^3+^ in the absence (**A**) and presence of kanamycin (**B** and **C**) for plasmid maintenance. Flasks were inoculated at an OD_600_ of 0.01 (**A**) or 0.03 (**B** and **C**) with washed cells from M9 overnight cultures grown with succinate in the absence (**A**) or presence (**B** and **C**) of kanamycin and 0.2% rhamnose as inducer for the plasmids pJEM[PedR2] and pJEM[PedR2^D53A^]. **A**) Growth of Δ*pedE* (black circles), Δ*pedE*_PedS2^S178P^ (orange dots), and Δ*pedE*_PedS2^S178P^Δ*pedR2* (grey dots). **B**) Growth of Δ*pedH*_PedS2^S178P^Δ*pedR2* harboring pJEM[PedR2] (grey dots) or pJEM[PedR2^D53A^] (orange dots). **C**) Growth of Δ*pedE*_PedS2^S178P^Δ*pedR2* harboring pJEM[PedR2] (grey dots) or pJEM[PedR2^D53A^] (orange dots). Data points represent the mean of biological triplets and error bars correspond to the respective standard deviations (positive error values).

### The conserved phosphorylation site D53 in PedR2 is essential for the REE-mediated switch

In order to study the essentiality of the phosphorylation site at position D53 of PedR2 (Bourret et al., 1990; Milani et al., 2005), we used inducible constructs for the production of the wild-type PedR2 protein (pJEM[PedR2]) and a mutated variant, in which the conserved aspartate in the CheY-like receiver domain was replaced by an alanine (pJEM[PedR2^S178P^]). After transformation of these plasmids into strains ∆*pedH*∆*pedR2* and ∆*pedE*_PedS2^S178P^ ∆*pedR2* their growth with 2-phenylethanol in the absence and presence of La^3+^ was monitored. When the plasmid-borne wild-type regulator PedR2 was induced in cells of ∆*pedH*_PedS2^S178P^ ∆*pedR2*, growth was observed after a lag-phase of < 24 h whereas the PedR2^D53A^ variant was unable to restore PedE-dependent growth in the same strain (Figure 4B). Intriguingly, the reverse result was obtained in strain ∆*pedE*_PedS2^S178P^ ∆*pedR2*. Here, the PedR2^D53A^ variant allowed PedH-dependent growth, whereas the wild-type regulator PedR2 did not lead to significant growth within 120 h of incubation (**Figure 4C**).

## DISCUSSION

We recently demonstrated that in *P. putida* KT2440 the production of the two PQQ-EDHs PedE and PedH is both tightly and inversely regulated depending on lanthanide availability, representing the first reported REE switch for PQQ-EDHs in a non-methylotrophic organism (Wehrmann et al., 2017). While we were able to show that Ln^3+^-dependent transcriptional activation of *pedH* is mostly, but not entirely, dependent on the presence of the PedH protein itself, the Ln^3+^-dependent transcriptional repression of *pedE* remained elusive. In this study, we present a detailed characterization of the mechanism underlying PedE and PedH regulation (Figure 5), in which the TCS PedS2/PedR2 acts as an essential signaling module for the REE-mediated switch between the two quinoproteins. Similar to the recently characterized spontaneous mutant of *M. buryatense* (Chu et al., 2016), we found that a single non-synonymous mutation within the periplasmic region of the sensor histidine kinase PedS2 (PedS2^S178P^), which differs from the LapD/MoxY domain found in MxaY of *M. buryatense* (Figure 1), is sufficient to terminate the Ln^3+^-mediated repression of *pedE*. Notably, various mutations at different sites of the protein can cause the observed suppressor phenotype. This might explain the repeatedly and fast occurrence of these suppressor mutants in our experiments and would support a similar notion in *M. buryatense* (Chu and Lidstrom, 2016; Chu et al., 2016).

**Figure 5:**
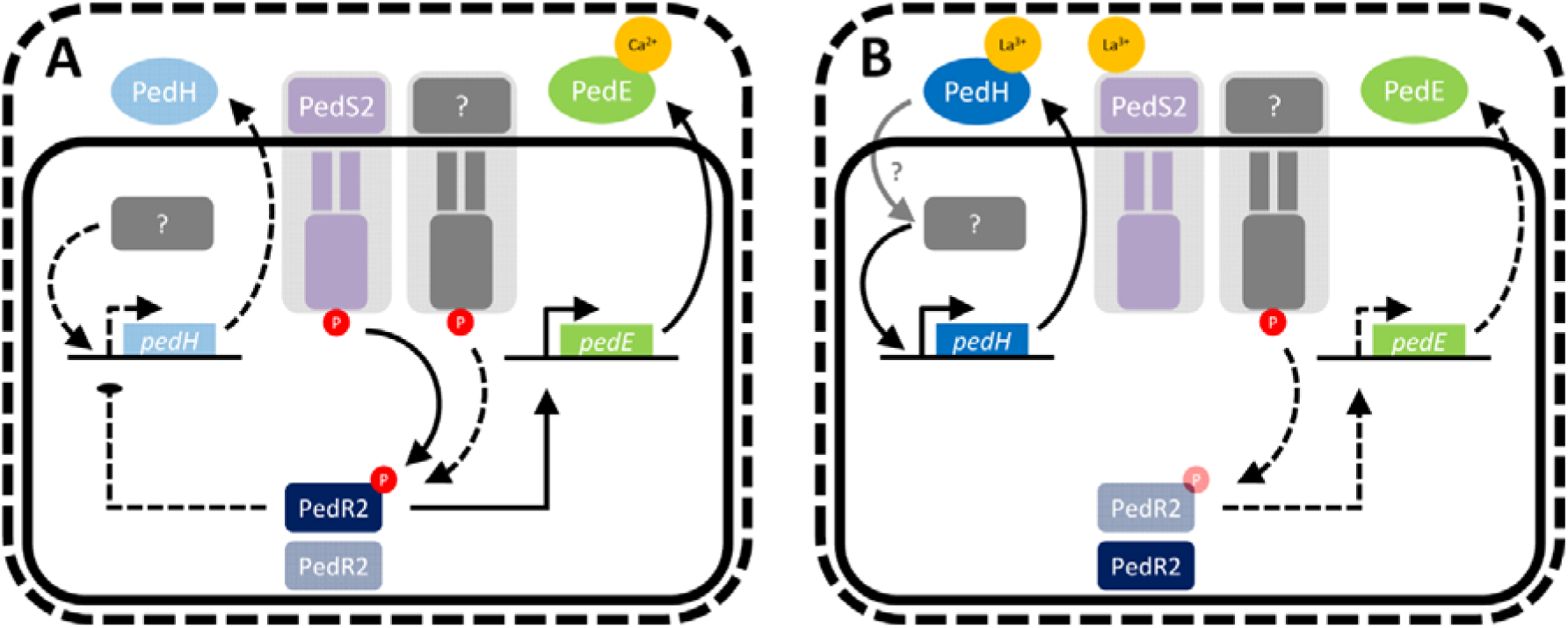
Working hypothesis of REE-mediated switch of *pedE* and *pedH* in *Pseudomonas putida* KT2440. Black lines indicate known regulations or functionalities. Grey boxes with question marks indicate unknown regulatory components. Solid lines indicate a strong regulatory effect on genes or production of enzymes, whereas dotted lines indicate weaker regulatory effects or low production of enzymes. A detailed description of the working model is given in the text.

Besides the essentiality for *pedE* regulation, our experimental data further provide strong evidence that PedS2 is also involved in the repression of *pedH* in the absence of lanthanides, but not in its Ln^3+^-dependent activation. This is based on the observation that the Δ*pedE*Δ*pedS2* deletion strain did not show any differences in growth and *pedH* regulation, while the suppressor mutant Δ*pedE*_PedS2^S178P^ displayed decreased *pedH* promoter activity and a strongly increased lag-phase in growth experiments in the presence of La^3+^. We additionally show that the *pedS2*-dependent regulations of *pedE* and *pedH* are both mediated by the LuxR-type response regulator PedR2. From these data we conclude that in absence of La^3+^ ions the PedS2 sensor histidine kinase is active and triggers phosphorylation of PedR2 at the conserved position D53 (PedR2^P^) (Bourret et al., 1990; Milani et al., 2005). The phosphorylated state of PedS2 subsequently has a dual regulatory function, namely the activation of *pedE* and concomitant repression of *pedH* transcription (**Figure 5A**). In this context, it is interesting to note that expression of the gene *exaA* in *Azospirillum brasilense* Sp7, which encodes for a PQQ-dependent alcohol dehydrogenase, is dependent on σ^54^ and its interaction with a LuxR-type response regulator that shares 45% sequence identity with PedR2 (Singh et al., 2017). Whether the transcriptional activation of *pedE* is dependent on a similar interaction of PedR2 with a specific sigma factor is currently unknown.

To our surprise, also a Δ*pedH*Δ*pedS2* mutant was able to grow on 2-phenylethanol, even though with an increased lag-phase and low *pedE* promoter activities (**Figure 2A3**). This indicates that at least one additional kinase beside PedS2 must be capable of phosphorylating PedR2 to facilitate growth of this strain under these conditions. Given that such an additional kinase exists, it is surprising that a *pedH* single mutant, in contrast to the aforementioned Δ*pedH*Δ*pedS2* double mutant, is only able to grow in the presence of REEs when PedS2 is mutated (e.g. PedS2^S178P^; **Figure 2A1**). This suggests that the activity of the additional, non-specific histidine kinase towards PedR2 in the presence of La^3+^ is repressed as long as PedS2 is functional. We hence propose that PedS2, as many other sensory histidine kinases in bacteria (Stock et al., 2000; Rowland and Deeds, 2014; Agrawal et al., 2016), also exhibits phosphatase activity on PedR2^P^ in the presence of La^3+^, thereby ensuring specificity of the signal transduction pathway and eliminating the interference from other non-specific kinases. In our working hypothesis, the presence of lanthanides in the medium leads to the repression of PedS2 kinase activity, most likely by direct binding of the metal ions to its periplasmic domain (**Figure 5B**). The reduced kinase- and postulated phosphatase activity of PedS2 consequently leads to the accumulation of unphosphorylated PedR2, which finally results in the loss of its regulatory functions. In addition, the transcription of *pedH* is activated *via* a yet unknown pathway, in which a functional PedH protein is an essential component most likely by acting as a lanthanide sensor (Wehrmann et al., 2017).

In conclusion, it appears that the REE-mediated switches in *M. extorquens* AM1 and *M. buryatense* during growth with methanol are predominantly dependent on only one lanthanide responsive pathway, which either proceeds via the XoxF1 and XoxF2 proteins or MxaY (Chu et al., 2016; Vu et al., 2016). Our results establish that in *P. putida* KT2440, a combination of at least two independent pathways are important to orchestrate the inverse regulation of *pedE* and *pedH* in response to lanthanides efficiently.

Several recent studies suggest that in methano- and methylotrophic bacteria, the REE switch might affect more genes than only those needed for the periplasmic oxidation system itself (Gu and Semrau, 2017; Good et al., 2018; Masuda et al., 2018). Reports on physiological consequences are, however, inconsistent as some studies found no effects (Nakagawa et al., 2012; Chu and Lidstrom, 2016; Vu et al., 2016; Masuda et al., 2018), whereas other studies reported a stimulating effect on biofilm formation, growth rates, and overall yields in the presence of REE (Fitriyanto et al., 2011b; Nakamura, et al., 2011; Good et al., 2018). We think it is not unlikely that additional REE-mediated regulatory effects do also exist in *P. putida* KT2440 in a context-dependent manner. Thus, one of our current foci is to investigate the global regulatory impact and physiological consequences of the presence and absence of lanthanides under varying environmental conditions.

## MATERIAL AND METHODS

### Bacterial strains, plasmids and culture conditions

A detailed description of the strains and plasmids used in this study can be found in **Table 1**. If not stated otherwise *Escherichia coli* and *P. putida* KT2440 strains were maintained on solidified LB medium. Routinely, strains were cultured in liquid LB medium (Maniatis et al., 1982) or a modified M9 salt medium (Wehrmann et al., 2017) supplemented with 25 mM succinate or 5 mM 2-phenylethanol as source of carbon and energy at 30°C and shaking, if not stated otherwise. For maintenance and selection, 40 µg mL^−1^ kanamycin or 15 µg mL^−1^ gentamycin for *E.coli* and 40 µg mL^−1^ kanamycin, 20 µg mL^−1^ 5-fluoro uracil, or 15 µg mL^−1^ gentamycin for *P. putida* strains was added to the medium, if indicated.

**Table 1:**
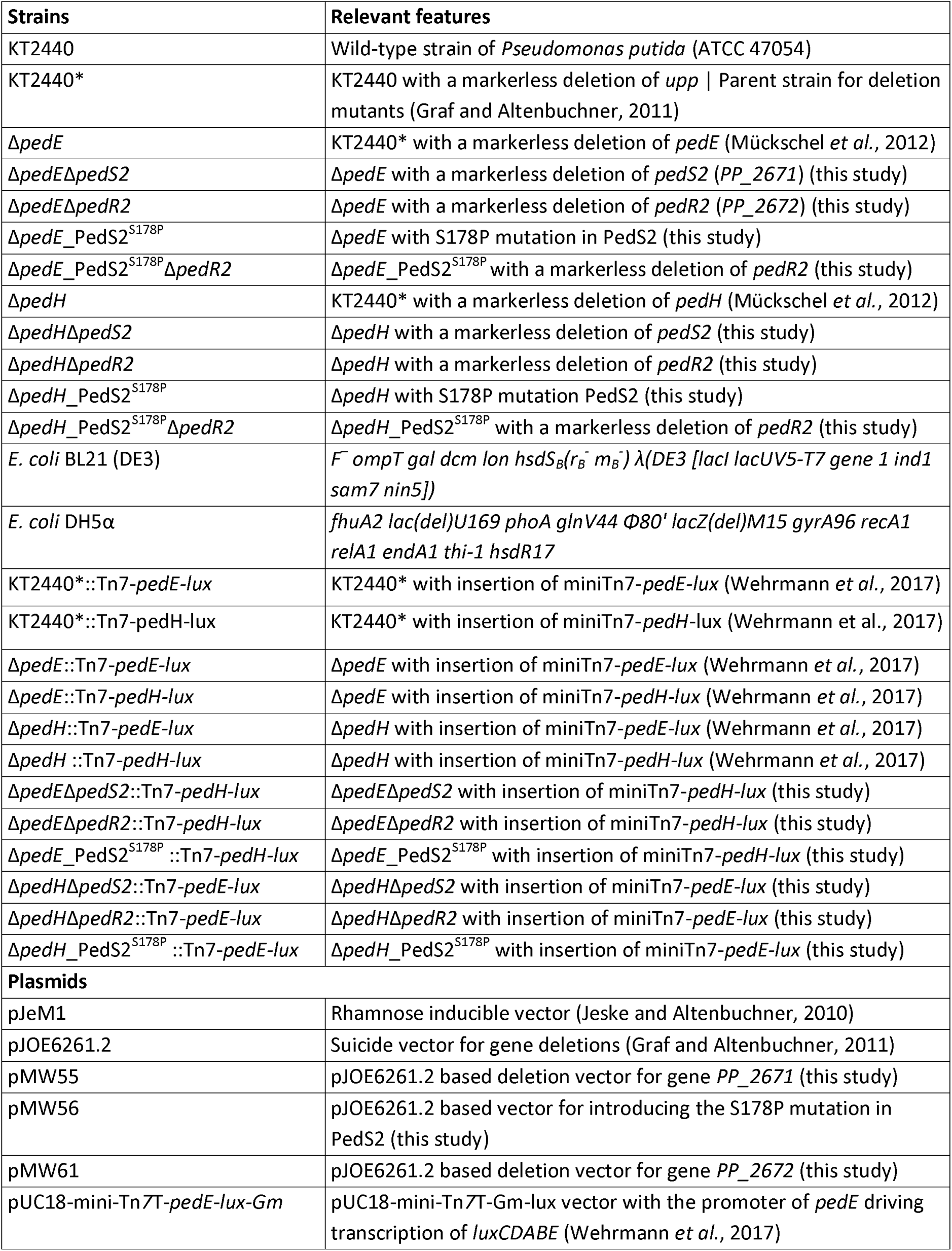
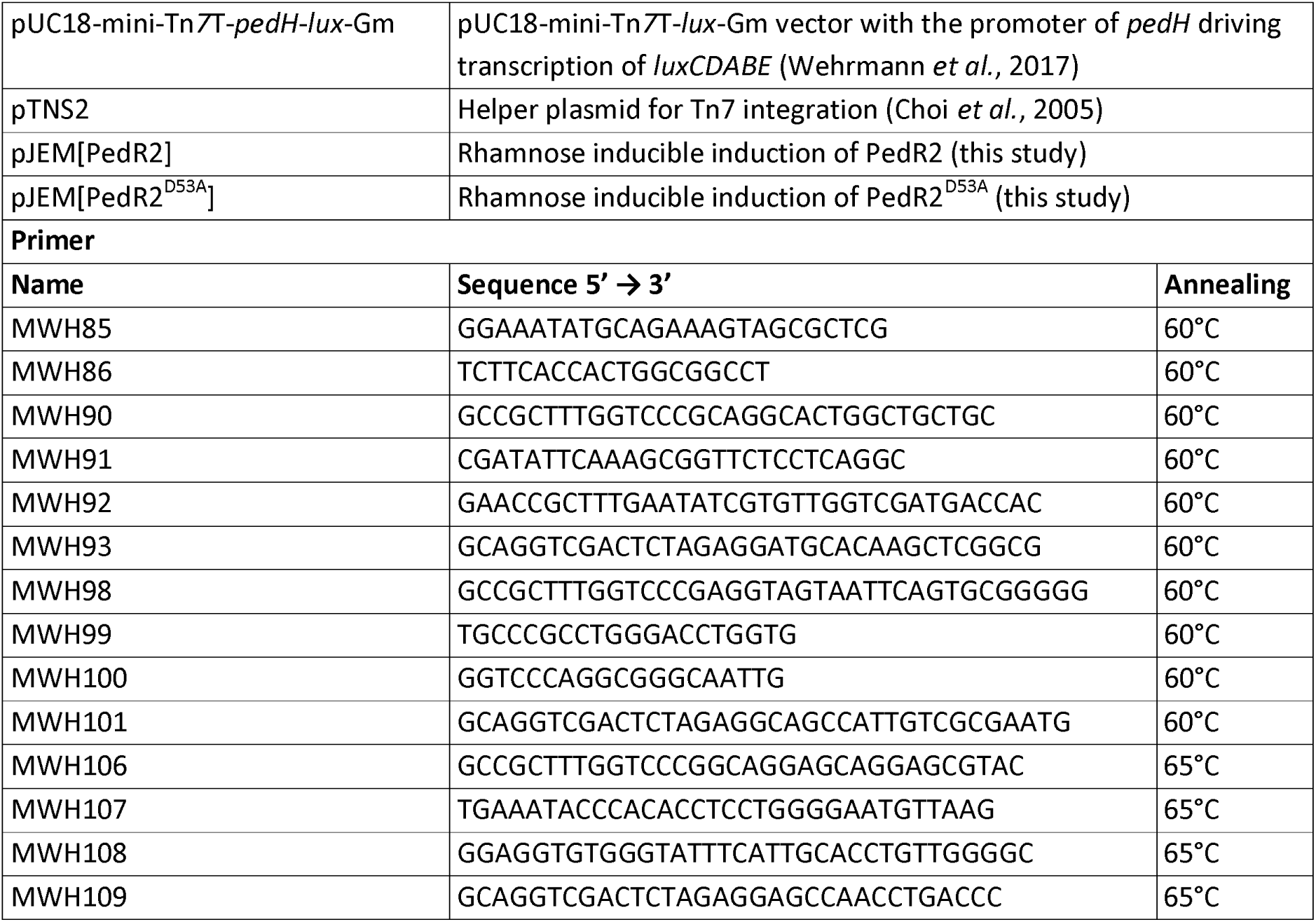
Strain, plasmids, and primer used in the study

### Liquid medium growth experiments

All liquid growth experiments were carried out using modified M9 with 25 mM succinate or 5 mM 2-phenylethanol as sole source of carbon and energy (see above) in 125 ml or 250 ml polycarbonate Erlenmeyer flask (Corning) or in 96-well 2 ml deep-well plates (Carl Roth), as described previously (Wehrmann et al., 2017). Briefly, washed cells from overnight cultures grown with succinate at 30°C and 180 rpm shaking were used to inoculate fresh medium with an OD_600_ of 0.01 and incubated at 30°C and 180 rpm (growth experiments in polycarbonate Erlenmeyer flasks) or 350 rpm (growth experiments in 96-well plates) shaking.

### Construction of plasmids

For construction of the deletion plasmids pMW55, pMW56 and pMW61 the 600 bp regions upstream and downstream of gene *pedS2* (*PP_2671*), amino acid residue S178 in gene *pedS2* (*PP_2671*), or gene *pedR2* (*PP_2672*) were amplified from genomic DNA of *P. putida* KT2440 using primers MWH90-93, MWH98-101 or MWH106-109 (**Table 1**). The two up- and downstream fragments and BamHI digested pJOE6261.2 were then joined together using one-step isothermal assembly (Gibson, 2011). Upon subsequent transformation of the constructs into *E. coli* BL21(DE3) cells, the correctness of the plasmids was confirmed by Sanger sequencing. Plasmids pJeM[pedR2] and pJEM[PedR2^D53A^] encoding for PedR2 or PedR2 with mutated amino acid residue 53 (D→A) under a rhamnose inducible promoter were ordered from an external source (Eurofins).

### Strain constructions and isolation of suppressor mutants

The *pedS2* (*PP_2671*) and *pedR2* (*PP_2672*) negative mutants as well as the PedS2^S178P^ allele were constructed using a recently described system for markerless gene deletion in *P. putida* KT2440 (Graf and Altenbuchner, 2011). Briefly, the integration vectors harboring the up- and downstream regions of the target genes (pMW55 and pMW61) or the up- and downstream regions of the region to be mutated including the desired S178P mutation (pMW56) were transformed into *P. putida* KT2440* and kanamycin (Kan) resistant and 5-flourouracil (5-FU) sensitive clones were selected on LB Kan agarose plates. After incubation at 30°C for 24 h in LB medium without selection markers, clones that were 5-FU^R^ and Kan^S^ were tested for successful gene deletion using primer pair MWH90/MWH93 or MWH106/MWH109 for gene *pedS2* or *pedR2*, respectively. The presence of the underlying PedS2^S178P^ mutation was verified by Sanger sequencing after gene *pedS2* of 5-FU^R^ and Kan^S^ clones was amplified using primer pair MWH85/MWH86. ∆*pedH* suppressor mutant strains were isolated from 25 ml liquid M9 cultures with 5 mM 2-phenylethanol as sole source of carbon and energy supplemented with 10 µM La^3+^ upon > 5 d incubation at 30°C and 180 rpm. Strains were passed thrice over LB agar plates and reevaluated for growth in liquid M9 medium supplemented with 5 mM 2-phenylethanol and 10 µM La^3+^. From cultures that showed growth upon a similar lag-phase as the KT2440* wild-type strain, the *pedS2* gene was amplified by PCR using primer pair M85/M86 and mutations were identified by Sanger sequencing.

To construct reporter strains for the analysis of *pedE* and *pedH* promoter activity in different genetic backgrounds, plasmids pUC18-mini-Tn*7*T-*pedE*-*lux-Gm* and pUC18-mini-Tn*7*T-*pedH*-*lux-Gm* (Wehrmann et al., 2017) were co-electroporated with the helper plasmid pTNS2 into selected mutant strains of *P. putida* KT2440 (**Table 1**). Proper chromosomal integration of the mini-Tn7 element in gentamycin resistant transformants was verified by colony PCR using P_put-_*_glm_*_SDN_ and P_Tn7R_ primers as described previously (Choi et al., 2005).

### Reporter gene fusion assays

For quantitative measurement of *pedE* and *pedH* promoter activity, strains of *P. putida* harboring a Tn*7*-based *pedE-lux* or *pedH-lux* transcriptional reporter fusion were grown overnight in LB medium with gentamycin (15 μg.ml^−1^), diluted to an OD_600_ = 0.2 in fresh LB medium and grown to an OD_600_ of 0.6. Cells were then washed three times in M9 medium without carbon source and finally adjusted to an OD_600_ of 0.2 in M9 medium with 1 mM 2-phenylethanol. For luminescence measurements, 198 µl of cell suspension was added to 2 µl of a 100 µM LaCl_3_ solution in white 96-well plates with a clear bottom (µClear; Greiner Bio-One). Microtiter plates were placed in a humid box to prevent evaporation and incubated at 28°C with continuous agitation (180 rpm), and light emission and OD_600_ were recorded at regular intervals in an FLX-Xenius plate reader (SAFAS, Monaco) for 6 h. For both parameters, the background provided by the M9 medium was subtracted and the luminescence was normalized to the corresponding OD_600_. Experiments were performed with biological triplicates, and data are presented as the mean value with error bars representing the corresponding standard deviation.

### Sequence identity determination

Protein sequence identities were determined based on amino acid sequence alignments of the proteins of interest generated using the Clustal Omega multiple sequence alignment tool (Sievers et al., 2014).

## FUNDING INFORMATION

The work of Matthias Wehrmann and Janosch Klebensberger was supported by an individual research grant from the Deutsche Forschungsgemeinschaft (DFG, KL 2340/2-1). The work of Charlotte Berthelot and Patrick Billard was supported in part by Labex Ressources21 (ANR-10-LABX-21-01).

## ACKNOWLEDGEMENTS

The authors would like to thank Prof. Bernhard Hauer for his continuous support. The authors further declare that the research was conducted in the absence of any commercial or financial relationships that could be construed as a potential conflict of interest.

